## Supplemental figures for "Locally released somatostatin triggers cAMP and Ca^2+^ signaling in primary cilia to modulate pancreatic β-cell function"

### Supplementary Data (Incedal-Nilsson et al)

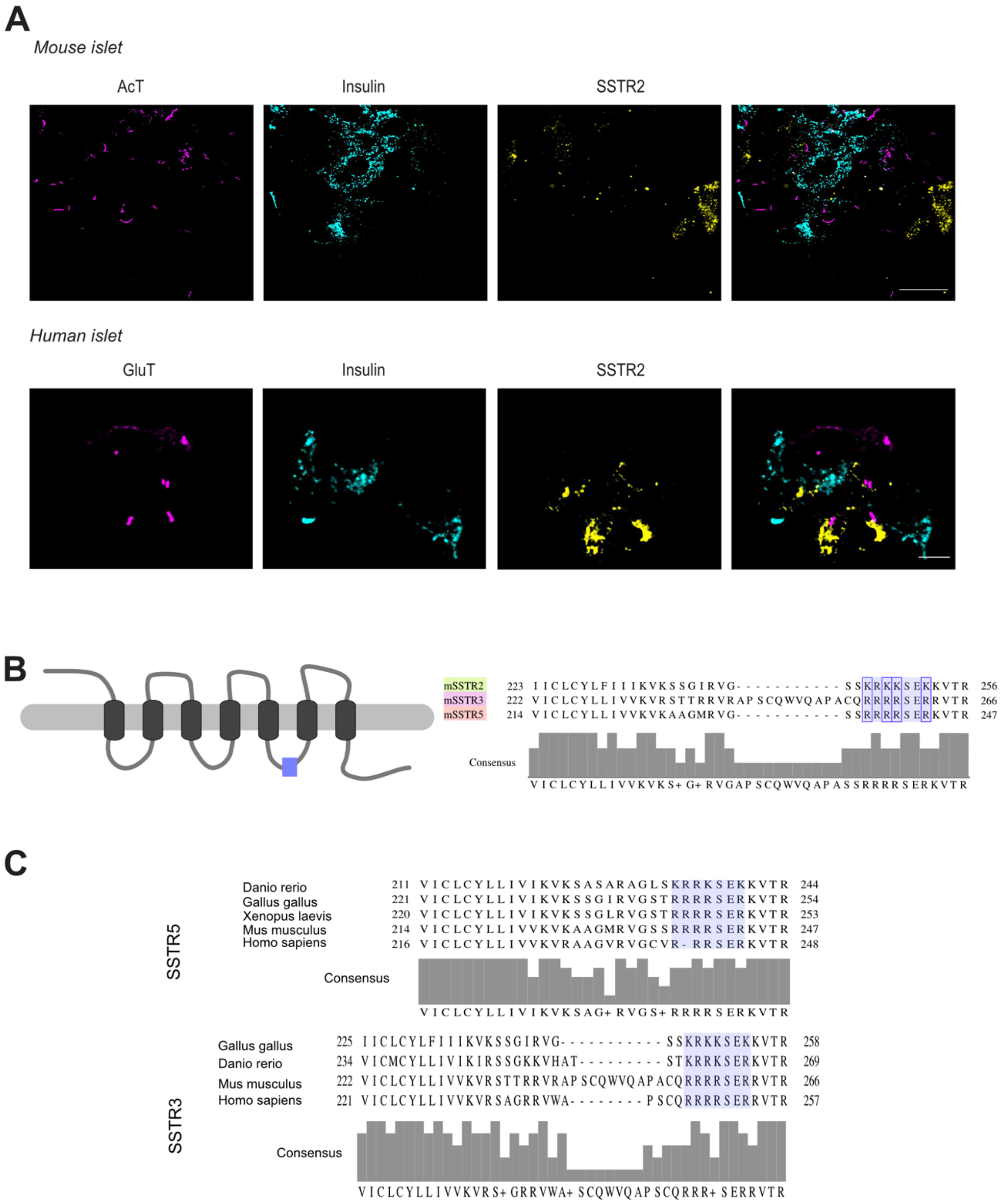


**Supplementary figure 1. A.** Confocal microscopy images of a mouse and human islet immunostained against SSTR2 (yellow), acetylated tubulin and glutamylated tubulin as cilia markers (magenta) and insulin (cyan).
**B.** An illustration shows the membrane localization of SSTRs, with IC3 loop is indicated by the purple box. Sequences of mSSTR2, mSSTR3 and mSSTR5. Conserved motif RxRxxR is highlighted.
**C.** Sequences of SSTR5 and SSTR3 showing the evolutionary conservation of a stretch of amino acids in purple.

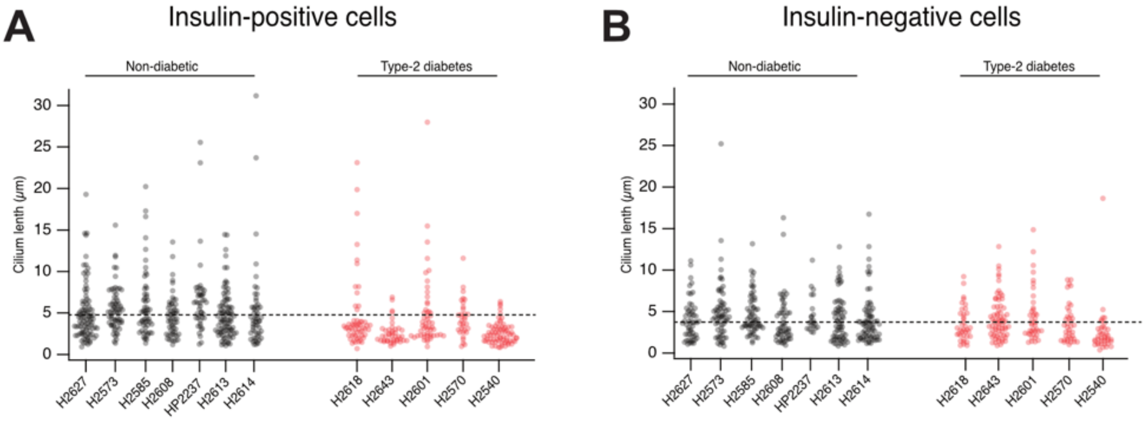


**Supplementary figure 2. A.** Quantification of cilium length in insulin-positive β-cells in islets from 7 non-diabetic (black) and 5 type-2 diabetic (red) human organ donors. Dashed line shows average for all non-diabetic donors.

**B.** Quantification of cilium length in insulin-negative cells in islets from 7 non-diabetic (black) and 5 type-2 diabetic (red) human organ donors. Dashed line shows average for all non-diabetic donors.

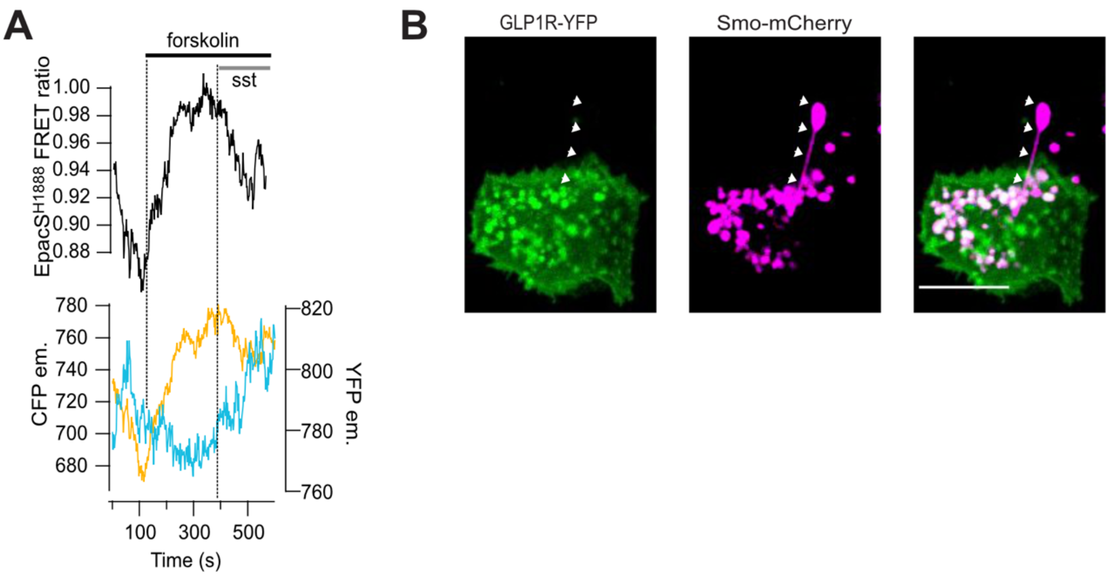


**Supplementary figure 3. A.** Representative recording of mArl13b-188 FRET ratio change from a MIN6 cell in response to 10 µM forskolin and 100 nM somatostatin.

**B.** Representative confocal microscopy images showing non-ciliary localization of GLP-1R (green) in MIN6 cells and mouse islets. MIN6 cells overexpress GLP-1R-YFP (green) and Smo-mCherry (magenta).

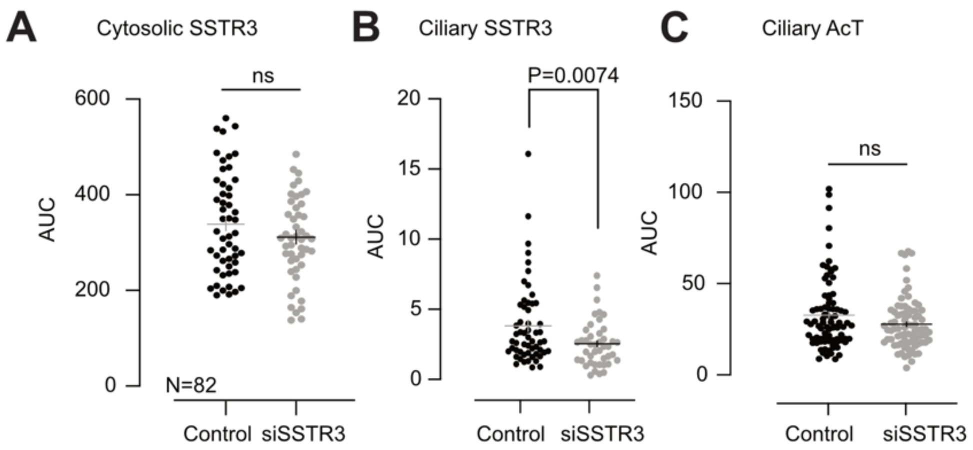


**Supplementary figure 4. A.** Quantification of cytosolic area selected in MIN6 pseudo islets. Fluorescence intensity is from control (black) and SSTR3KD (gray) cells immunostained for SSTR3. (SEM±, $n_{ctrl}$=50 and $n_{KD}$=48, 3 different preparations, no statistical difference by Mann-Whitney t-test, unpaired.)

**B, C.** Quantifications of line profiles drawn along cilia of MIN6 pseudo islets for SSTR3 and acetylated tubulin. Notice that SSTR3 signal is reduced in cells treated with SSTR3 siRNA on the right, and acetylated tubulin signal is unaffected on the left. (p value is 0.0074 for SSTR3 comparison by Mann-Whitney, $n_{ctrl}$=83 and $n_{KD}$=86, 3 different preparations).

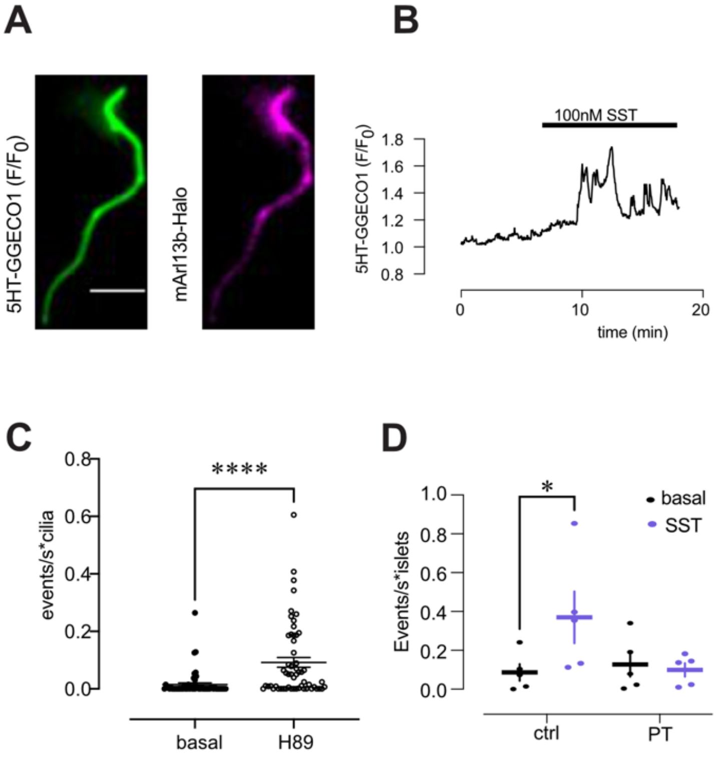


**Supplementary figure 5. A.** A TIRF microscopy image from a mouse islet expressing 5HT6-GGECO1 and mArl13b-Halo from the insulin promoter.
**B.** Representative trace from an identified β-cell of a mouse islet showing ciliary $Ca^2$ in response to 100 nM somatostatin.
**C.** Event count of ciliary $Ca^{2+}$ changes from mouse islets in response to 10 µM H89 (7 islets; 56 cilia; 3 different preparations. ****p<0.0001, Wilcoxon matched pair t-test).
**D.** MIN6 pseudo-islets were cultured under control condition (black) or in the presence of pertussis toxin (purple) for 18h. Quantifications of the $Ca^{2+}$ responses to 100 nM somatostatin are shown to the right ($n_{ctrl}$=5, $n_{PT}$=5 islets from 3 different preparations, SST response in control p=0.0101, SST response in PT not significant, assessed by Sidak´s multiple comparison).

**Table 1 – Donor characteristics**

| | **Non-diabetic** | **Type-2 diabetic** |
|---|---|---|
| Gender | 6 male/0 female | 4 male/1 female |
| Age | 60±6 years (50-69) | 73±6 years (68-80) |
| BMI | 28±4 kg/m$^2$ (22-33) | 29±2 kg/m$^2$ (28-32) |
| HbA1c | 37±3 mmol/mol (33-41) | 44±6 mmol/mol (37-48) |
| Cold ischemic time | 11h20min (7h59min-16h50min) | 12h40min (7h10min-20h57min) |
